# A confound-free method to manipulate pupil size in psychological experiments

**DOI:** 10.1101/2025.06.17.660124

**Authors:** Joshua Snell, Luise Wagner, Ana Vilotijević

## Abstract

Those who study visual perception and visuo-spatial attention are increasingly paying interest to the ways in which these processes are influenced by the size of the pupil. However, this realm of research is complicated by the pupil’s notorious susceptibility to confounders and difficulties in disentangling cause and effect. Recent studies have sought a solution in the exploitation of intrinsically photosensitive retinal ganglion cells (ipRGCs), which trigger pupil constrictions upon detecting blue light. Experiments typically present stimuli against blue versus red backgrounds of equal perceived luminance, to effectuate systematically smaller pupils in the former than in the latter condition. Here we provide a further validation of this method, by testing a scenario of potential concern. Via retino-hypothalamic pathways, ipRGC activation modulates alertness and arousal. Blue and red backgrounds may therefore differentially impact behavior in addition to the pupil, potentially confounding inferences about the pupil’s impact on cognition. This was investigated with an auditory task in which participants responded as quickly as possible to sequences of randomly timed beeps while looking at blue versus red displays. Each participant was tested with an easy and a difficult version of the auditory task. Results suggest that the ipRGC method is good to go: in the presence of effects of task difficulty on both pupil size and task performance, the display color exclusively influenced pupil size without affecting task performance.

## Introduction

The size of the pupil impacts visual processing in various ways: pupil constrictions (i) enhance foveal acuity, (ii) increase depth of field and (iii) enhance foveal vision through reducing light scatter; but also (iv) reduce peripheral awareness and (v) impair vision in low-light conditions. Pupil dilations necessarily do the precise opposite. But beyond these basic sensory consequences, might the size of the pupil also affect certain cognitive processes, such as attention or memory? Whereas the pupil has long been used as a read-out of various cognitive processes (e.g., *arousal*: Joshi et al. 2016; Joshi and Gold 2020; Murphy et al. 2014, *mental effort*: Granholm et al., 1996; Hess & Polt, 1960; Kahneman & Beatty, 1966, *attention*: Binda et al., 2014; Binda & Murray, 2015; Bombeke et al., 2016; Mathôt et al., 2013; Naber & Nakayama, 2013; Unsworth & Robison, 2017; Snell et al., 2018; Vilotijević and Mathot, 2023, Vilotijević and Mathot, 2024, Vilotijević and Mathôt, 2025), cognitive neuroscience has only recently begun to consider the pupil’s potential role as an effector in its own right (e.g., Suzuki et al., 2019; Mathôt et al., 2023; Vilotijević & Mathôt, 2024). This, however, begs ways to disentangle cause and effect; and in the case of the pupil this happens to be notoriously difficult. Consider, for example, visuo-spatial attention. When attention is more widespread and/or shifts towards the peripheral visual field, the pupils tend to dilate (e.g., Vilotijević & Mathôt, 2023). With larger pupils, there is more retinal illumination, particularly in the periphery. This enhances peripheral visual processing, which increases the potential of peripheral stimuli to capture attention; and with the resulting attentional shift to the peripheral visual field we have come full circle. Various such ‘loops’ exist between the pupil and other cognitive processes (for a review, see Mathôt, 2018), and scientists have yet to pinpoint the causal relationships.

How to solve the above conundrum? Intriguingly, recent studies suggest that the solution may be as simple as presenting stimuli on red versus blue backgrounds. In addition to the more well-known rods and cones, the retina comprises intrinsically photosensitive retinal ganglion cells (ipRGCs; e.g., Berson, 2003; McDougal & Gamlin, 2010; Spitschan et al. 2017; Zele et al. 2018). These do not directly project onto the visual cortex, meaning they do not engender conscious visual percepts. Instead, they are mainly involved in non-image-forming functions such as the regulation of circadian rhythms and pupillary control, through their projections onto the suprachiasmatic nucleus (SCN) of the hypothalamus and the olivary pretectal nucleus (OPN) of the midbrain, respectively (e.g., Berson, 2003; McDougal & Gamlin, 2010; Ecker et al., 2010; Fan et al., 2018). ipRGCs contain the photopigment melanopsin, which has peak sensitivity to blue light (∼480 nm). Their activation brings about a relatively slow (onsetting at ∼5s) but sustained pupillary constriction. As such, it would seem that the size of the pupil can be very easily and non-invasively manipulated by presenting participants with isoluminant red versus blue displays, for respectively larger versus smaller pupils (e.g., Wardhani et al., 2022; Kinzuka et al., 2022; Mathôt et al., 2023; Vilotijević & Mathôt, 2024).

The ipRGC method has various benefits compared to other methodologies. For instance, manipulation of pupil size through overall display luminance (e.g., presenting black versus white backgrounds) potentially introduces differences in the visibility of stimuli or different degrees of physical strain on the eyes. These problems are avoided with the ipRGC method because the red and blue displays are (perceptually) isoluminant. Alternatively, pupil size may be manipulated pharmacologically, e.g. by instilling a homatropine hydrobromide solution in the eye’s conjunctival sac (e.g., Campbell & Gregory, 1960); but this may be accompanied by blurred vision (since the eye’s ability to accommodate is impaired) and differential behavior because of discomfort and physical sensations during and after the instillation.

That being said, the ipRGC method has only been used in a handful of studies and warrants further validation (Mathôt et al., 2023; Wardhani et al. 2022). A potential point of concern is the retino-hypothalamic pathway: upon activation, the SCN—through pathways involving the locus coeruleus (LC) (Vandewalle et al., 2007), can enhance alertness and arousal. As such, there is a possibility that the ipRGC method inadvertently impacts behavior, with, e.g., better task performance with blue backgrounds relative to red backgrounds. Note, there is at least one reason to believe that the SCN’s role is relatively minor compared to that of the retino-tectal pathway and the OPN: arousal is associated with dilated pupils (e.g. Bradley et al., 2008; Mathôt, 2018), meaning that, as far as the SCN is concerned, the pupil should be larger in blue light—but this is evidently not the case (Wardhani et al., 2022; Kinzuka et al., 2022; Mathôt et al., 2023; Vilotijević & Mathôt, 2024). Yet, even if the OPN has a firmer grip on the pupil than the SCN, the latter may nevertheless separately exert an inadvertent influence on behavior, and this would pose a problem to the ipRGC method. With the present study we hope to rule out this scenario.

### The present study

Here we investigated whether ipRGC activation impacts behavior in a response time task. Participants responded as quickly as possible to auditory cues, while looking at red versus blue displays. Participants were tested in both an easy and a difficult version of the auditory task. We expected that the difficult task would provoke more mental effort and arousal, and that this would affect the pupil concurrently with task performance. Against this reference variable, we were hoping to see that the color of the display would exclusively affect the pupil without impacting task performance.

## Methods

### Participants

Thirty-eight students (age M=21.0, SD = 3.0) from the Vrije Universiteit Amsterdam gave informed consent to participate in this study for course credit or monetary compensation. Participants reported to have no visual impairments or mental disabilities. Recruitment and data collection were done in full accordance with the Declaration of Helsinki. Ethical approval was obtained from the Ethics committee of the Faculty of Behavioural and Movement Sciences (approval code VCWE- 2021-009).

### Stimuli and design

Participants performed an auditory detection task while looking at red and blue displays. We used a 2×2 experimental design with task difficulty (*easy* versus *difficult*) and display color (*red* versus *blue*) as factors. The experiment consisted of four blocks—two easy (E) and two difficult (D)— and these were offered in a counterbalanced subset of block orders (i.e., we used EDED, DEDE, EDDE and DEED, but no EEDD or DDEE). The two display colors were used in the two respective halves of each block, again with counterbalanced order.

In the easy blocks, participants responded as quickly as possible with the down arrow key to a total of ninety-two 50 ms 440 Hz beeps played at ∼60 dB (46 with a blue display, 46 with a red display). Two beeps were played per 5s interval, with the timing between beeps randomly varying between 300 and 4700 ms. This means that if the second beep was played, say, 1750 ms after the first beep, then the next beep occurred 5000 - 1750 = 3250 ms after that (hence ensuring two beeps per 5s). In the difficult blocks, there were forty 440 Hz beeps and forty 458 Hz beeps. Here, participants had to respond with the up and down arrow keys to the higher- and lower- pitch beeps, respectively. In the difficult blocks, there were also twelve high-pitch (660 Hz) double-beeps (two 40 ms tones separated by 55 ms silence). Upon hearing a double-beep, participants had to refrain from pressing any key. The total of ninety-two beeps was presented in random order.

For our displays, we adopted the luminosities that were established by Mathôt et al. (2023) to have equal subjective intensity, at 6.74 and 9.84 cd/m^2^ for red and blue, respectively. The equal subjective intensity was verified by the fact that the initial pupillary light response, as triggered by rod and cone cells in the first ∼2.5s (which is earlier than the ipRGC-driven sustained pupil constrictions), differed non-significantly between the red and blue displays.

### Procedure

Participants were seated in a dimly-lit room where we provided task instructions orally and onscreen. Participants were instructed to focus at a central fixation dot throughout the experiment. The eye position and pupil size were recorded with an EyeLink 1000 eye-tracker (SR Research, Canada). The experiment was implemented in OpenSesame (Mathôt et al., 2012) with the PyGaze package for eye-tracking (Dalmaijer et al., 2014). At the start of each block, participants were instructed whether they were about to do the easy or difficult task. Each block started with a 7s interval during which we showed the display in red or blue without presenting any beeps. These were so-called *inducer* intervals (e.g., Mathôt et al., 2023), which served to trigger a difference in the sustained pupil response due to differential ipRGC activation. Then 46 beeps occurred (these could be regarded as 46 trials or data points). After every beep, participants had until the next beep to respond. After 46 beeps, the display changed to the other color, and after another 7s inducer, the remaining 46 beeps were presented. The total of 368 experimental trials was preceded by 10 practice trials with a gray display. The entire experiment took approximately 35 minutes.

### Pupil data processing

Following the workflow for preprocessing pupillary data (Mathôt & Vilotijević, 2022), we first interpolated blinks and downsampled the data by a factor of 10. Next, we baseline-corrected the data by subtracting the mean pupil size during the first 50 ms after the onset of the display from all subsequent pupil-size measurements on a trial-by-trial basis. Baseline correction was carried out separately for each combination of task difficulty (easy vs. difficult) and display color (red vs. blue). Finally, trials in which the baseline pupil size deviated by more than ±2 z-scores from the mean (6.25%) were considered outliers and excluded from further analysis.

## Results

Pupil size, response time (RT) and accuracy were analyzed with respectively linear mixed-effect models (LMMs) with task difficulty and display color as factors and participants as random effect. Models successfully converged with inclusion of the random slope for display color. We report *b*- values, standard errors (SEs) and *t*-values (RTs) or *z*-values (accuracy), with values | *b* | and | *z* | > 1.96 considered significant. Before all analyses, we excluded trials with RTs < 100 ms or > 1499 ms (∼8%), so that 12,568 data points remained.

As noted above, every beep was considered a trial. For our analysis of pupil size, we logged the pupil size at the time of responding. Although the analysis is based on one pupil size measurement per trial, we do report continuous pupil size data per block type in Figure 1.

**Figure 1.**
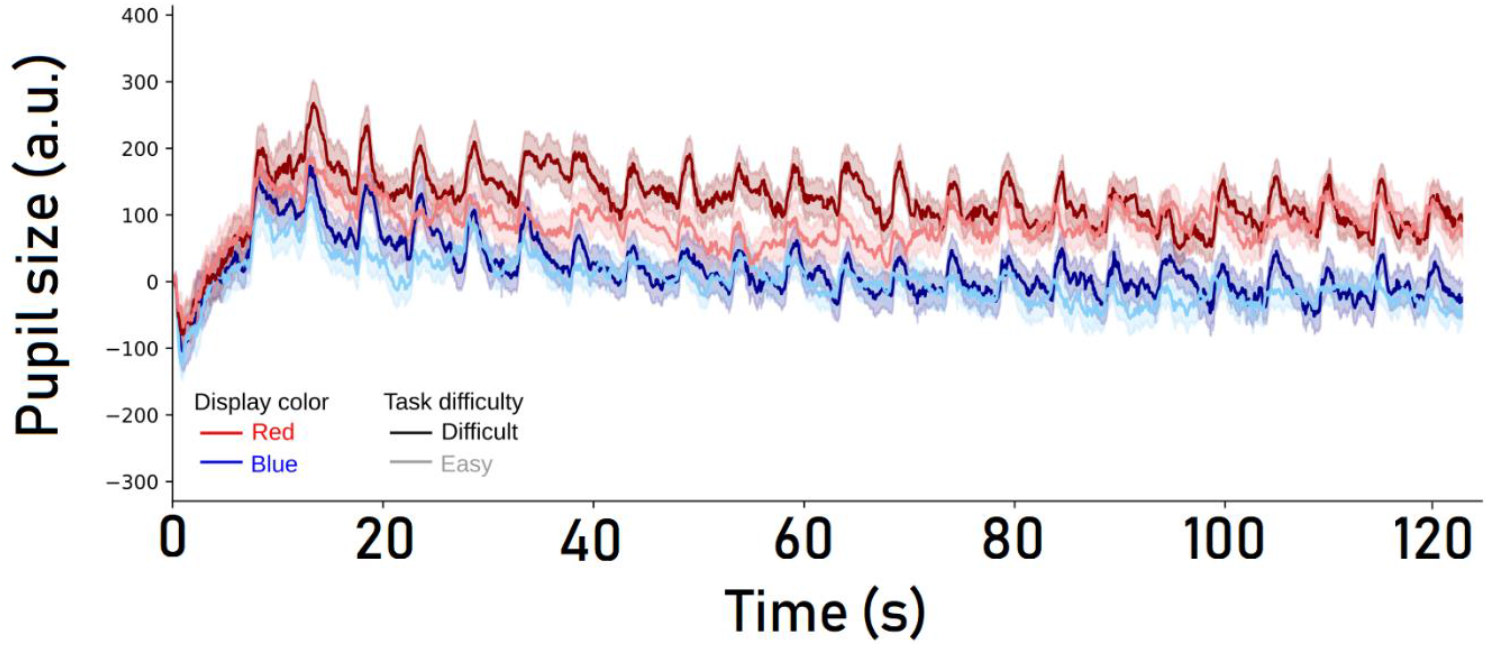
Baseline-corrected pupil size as a function of task difficulty (*easy* vs. *difficult*) and display color (*red* vs. *blue*), measured across 46 beeps (i.e., half a block). The shaded areas around each line reflect SEs. Abbreviations: a.u., arbitrary units.

In line with previous applications of the ipRGC method, blue displays engendered systematically smaller pupils than red displays (*b* = -53.27, SE = 21.97 *t* = -2.43). The pupil was also impacted by task difficulty, with smaller pupils in the easy blocks than in the difficult blocks (*b* = -20.99, SE = 4.82, *t* = -4.35). We also observed an interaction, such that the impact of task difficulty on pupil size was significantly greater with red than blue displays (*b* = 37.18, SE = 9.64, *t* = 3.86; Figure 2). We reflect on this interaction in the Discussion.

**Figure 2.**
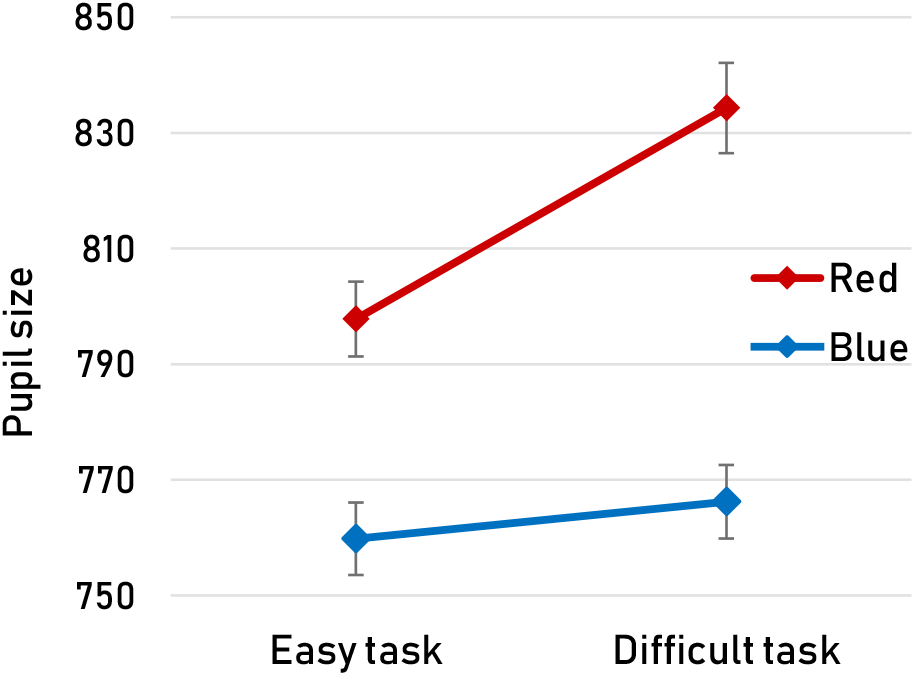
Overall average pupil size (arbitrary units, not baseline-corrected) as a function of task difficulty (*easy* vs. *difficult*) and display color (*red* vs. *blue*). Error bars represent SEs.

The effect of task difficulty on pupil size was accompanied by an effect in task performance, with slower responses and more errors in the difficult blocks (RT: *b* = 263.68, SE = 3.60, *t* = 73.16; accuracy: *b* = 2.69, SE = 0.07, *z* = 41.10). Such relations between behavior and pupil size may be considered common and unsurprising (e.g., Mathôt, 2018); but meanwhile, in the present study we were also hoping to find that the ipRGC-induced pupillary responses were *not* accompanied by behavioral differences. These hopes were fulfilled: in the presence of a clear effect of display color on pupil size, the display color did not affect task performance whatsoever (RT: *b* = 0.52, SE = 5.29, *t* = 0.10; accuracy: *b* = 0.03, SE = 0.06, *z* = 0.50; Figure 3).

**Figure 3.**
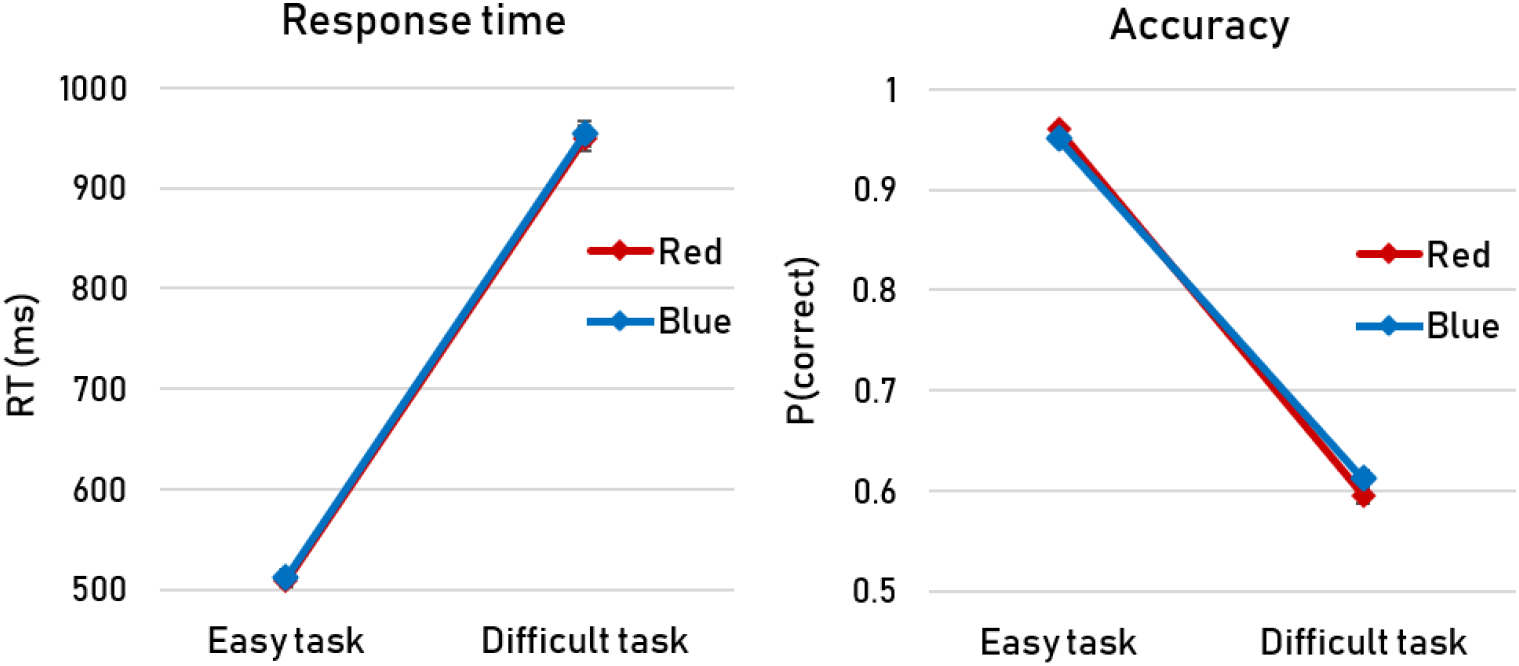
Average RTs and accuracies as a function of task difficulty (*easy* vs. *difficult*) and display color (*red* vs. *blue*). Error bars—which are so small that they are largely obscured by the value markers—represent SEs.

## Discussion

How does the size of the pupil affect various cognitive processes? This is an open problem, primarily because pupil size is determined by—and likely reciprocally determines—a myriad of factors. An important first step in solving this puzzle is the development of a technique to manipulate the pupil in a clean, confound-free manner. Various recent studies have championed the stimulation of intrinsically photosensitive retinal ganglion cells (ipRGCs) as the solution (e.g., Wardhani et al., 2022; Kinzuka et al., 2022; Mathôt et al., 2023; Vilotijević & Mathôt, 2024). The ipRGC approach is promising because ipRGCs themselves do not project directly onto the visual cortex (meaning they do not contribute directly to visual percepts), and because their exploitation—which involves the presentation of red versus blue displays—is strikingly simple and non-invasive.

The present study constitutes an important validation step. In addition to the retino-tectal pathway that underlies the pupillary response, ipRGCs project to the suprachiasmatic nucleus (SCN) and locus coeruleus (LC), which are involved in regulating alertness and arousal (e.g., Berson, 2003; Fan et al., 2018). Therefore, we needed to rule out that red and blue displays inadvertently impact behavior in addition to the pupil.

The ipRGC method has passed our validation test: in the presence of an impact of task difficulty on both pupil size and behavior, ipRGC stimulation solely affected pupil size without concurrently impacting behavior. We surmise that, within the scope of a typical cognitive psychology experiment, activation of the SCN-LC pathway is too minor to have a meaningful effect on behavior. This stands in sharp contrast to the retino-tectal pathway, the involvement of which is prominent enough to effectuate clear pupil size differences.

Although the present results are encouraging, they also provide a theoretical insight that is worth taking into account when employing the ipRGC method. We found that the modulation of pupil size by task difficulty was only observed in the presence of a red display. When participants viewed the blue display, pupil size remained stably small, regardless of task difficulty. This finding might suggest that the relationship between pupil size and arousal—long exploited in the study of arousal and/or mental effort (e.g., van der Wel & van Steenbergen, 2018; Kahneman, & Beatty, 1966; Klingner, et al., 2011; Hess, & Polt, 1964)—may be contextually constrained by sensory inputs that engage low-level photoreceptive pathways. One possibility is that stimulation of ipRGCs by blue light gates arousal-related influences on pupil size. The parasympathetically driven pupillary constriction would as such dominate, leaving little room for arousal-driven pupil dilations. Though speculative at this point, one consequence would be that one cannot simultaneously control pupil size via ipRGCs and read-out certain cognitive processes (e.g., mental effort, arousal) via pupil size modulations. Such processes are then better tracked via behavioral measures or, e.g., skin conductance. We should also emphasize that this does not undermine the ipRGC method in itself: after all, the ipRGC method’s main value lies in the ability to investigate the pupil’s influence on cognition, rather than the other way around.

On a final methodological note, might the story have been different if we had used a different behavioral task? Specifically, is there a possibility that we would observe a direct impact of ipRGCs on behavior if the task were more sensitive to small fluctuations in alertness and/or arousal? We reckon this is unlikely. Our task already appears to have been quite sensitive, in the sense that relatively small task-induced fluctuations in pupil size coincided with drastic fluctuations in behavior (Figure 3). By comparison, the ipRGC-induced pupil size effect was approximately twice as large (Figure 2), but there was clearly no impact on behavior whatsoever. It may furthermore seem reasonable to predict that things would look differently if our task had been visual rather than auditory in nature. But this is precisely the point: our auditory task shows that any impact of ipRGCs in a typical cognitive experiment (involving e.g. a detection or discrimination task) on behavior, if it were to exist at all, is negligible. Therefore, if any effect of ipRGC stimulation on behavior were to emerge in a visual task, researchers can safely draw a causal link to the size of the pupil rather than arousal.

In closing, here we have established that the ipRGC method is a clean and confound-free way to manipulate the size of the pupil. Furthermore, our data lead to believe that arousal-related dilations are not uniformly expressed, but are instead modulated by background and environmental illumination. We anticipate that the ipRGC method will have a prominent role in the ongoing endeavor to unravel the pupil’s impact on cognition.

## Acknowledgements

This work was supported by the European Research Council (grant ERC101164084 awarded to J.S). We thank Lesly Badtke, Simona Szabóova, Tamara Rolović, Marion Silliard, Miriam Kriegbaum and Danielle Lim for their assistance in piloting this experiment.

